# Acute sildenafil administration reduces susceptibility to induced atrial fibrillation in sheep

**DOI:** 10.1101/2024.09.09.612118

**Authors:** Nathan C Denham, George W P Madders, D C Hutchings, C E R Smith, A S Whitley, M Obeidat, A W Trafford, C M Pearman, K M Dibb

## Abstract

**Background:** Sildenafil is a PDE5 inhibitor with a very good safety profile and animal models suggest it may be beneficial in the treatment of heart failure and ventricular fibrillation. Sildenafil has also been associated with a reduced incidence of atrial fibrillation (AF) in a retrospective observational study. We have therefore sought to determine whether sildenafil has a direct effect on atrial electrophysiology and resultant AF burden.

**Methods:** Invasive electrophysiological studies were performed in 12 anaesthetised healthy adult female Welsh mountain sheep. Pacing protocols were performed in the right atrium before and after administration of an acute 10 mg intravenous bolus of sildenafil and the burden of AF assessed.

**Results:** Sildenafil profoundly reduced the vulnerability to AF, decreasing AF duration (112.2 ± 73.5 s vs. 3.3 ± 1.4 s), the number of burst pacing inductions causing AF (90 % vs 70 %) and the complexity of AF. The antiarrhythmic effects of sildenafil were determined to be resultant of prolongation of both the atrial effective refractory period (146.9 ± 7.2 ms vs 166.2 ± 32.5 ms) and the atrial excitation wavelength (12.9 ± 0.07 cm vs 15.0 ± 0.07 cm) and resulted in a shallower restitution curve, reflected in a decreased magnitude of monophasic action potential alternans (0.09 ± 0.001 mV vs 0.05 ± 0.10 mV).

**Conclusions:** In the subjectively healthy atria of a highly translational model a strong antiarrhythmic effect upon acute sildenafil application was observed suggestive of a potential clinical benefit in AF.

## Introduction

Atrial fibrillation (AF) needs novel therapeutic targets and treatment options. Use of the phosphodiesterase 5 inhibitor (PDE5i) sildenafil has been associated with a reduced incidence of AF in a retrospective observational study [1]. Sildenafil has also been reported to decrease spontaneous firing in rabbit pulmonary vein sleeves [2], but little is understood about its effects on atrial electrophysiology e.g. the atrial effective refractory period (AERP) or conduction velocity and it remains to be directly tested if sildenafil can decrease AF.

In the ventricle sildenafil is anti-arrhythmic at the cellular level, where acute application decreases ventricular arrhythmias in the rodent heart post-infarct, the canine heart during ischemia and in a sheep model of drug-induced long QT [3–5]. The mechanism has been shown to be mediated by a reduced L-type calcium current (*I_Ca-L_*) and consequent decrease in sarcoplasmic reticulum (SR) calcium content [5–10], as well as a blunted response to β-adrenergic stimulation [11–19].

The aim of this study was to investigate whether acute PDE5 inhibition with sildenafil could decrease the vulnerability to AF *in vivo* in a large animal model and determine the electrophysiological mechanism of any anti-arrhythmic effect in the atrium.

## Methods

All procedures accorded with The UK Animals (Scientific Procedures) Act (1986) and EU directive 2010/63. Institutional approval was received from The University of Manchester Animal Welfare and Ethical Review Board. Reporting follows the ARRIVE (Animal Research: Reporting of in vivo Experiments) guidelines.

### Baseline assessment

Twelve young adult (≈18 months) female Welsh Mountain sheep underwent baseline echocardiography (Vivid 7; GE Healthcare UK) where left atrial diameter was measured in the anteroposterior axis and averaged over three cycles. Sheep were then anaesthetised with 3% v/v isoflurane and underwent a baseline 5-lead electrocardiogram (ECG) (Emka Technologies, Paris, France) recorded continuously for one minute. Mean ECG data was analysed over the minute in bins of ten consecutive sinus beats using LabChart version 7 (AD instruments, Colorado Springs, CO) and included: resting heart rate (expressed at cycle length), PR interval, QRS duration, QT interval (uncorrected) and the maximum P wave duration in the lead with the best signal to noise ratio (PWD_max_).

### Invasive left ventricular pressure monitoring

Right carotid arterial access was obtained using The Seldinger technique and a 5Fr diagnostic coronary catheter (Cordis, Johnson and Johnson, USA) passed retrograde aortic into the left ventricle. Left ventricular systolic pressure (LVSP) and end diastolic pressures (LVEDP) were recorded simultaneously with the baseline ECG in LabChart. Prior to each recording, the transducer underwent a two-point calibration at zero and 200 mmHg (reported accuracy ±1 mmHg). Mean left ventricular pressures were measured over ten consecutive beats.

### Electrophysiology studies

Right jugular venous access was gained via surgical cut down and venotomy. Permanent endocardial pacing leads were advanced to the right ventricular apex and the right atrial appendage, after which they were connected to a dual chamber internal cardiac defibrillator (ICD; Medtronic, Woburn, MA).

Monophasic action potentials (MAPs) were recorded from the posterior wall of the right atrium using a Blazer electrophysiology catheter (Boston Scientific, Quincy, MA). The AERP was determined using programmed electrical stimulation through the ICD at basic cycle length 400 ms for eight beats followed by an extrastimulus decreasing in 10 ms decrements from 350 ms to 100 ms. The AERP was measured as the longest cycle length of the extrastimulus which failed to result in an atrial MAP. The MAP duration at 90 % repolarisation (APD90) was calculated using a custom program written in Microsoft Excel. Atrial MAP alternans was determined by fixed rate burst pacing through the ICD over a range of cycle lengths from 500 to 250 ms. The presence of MAP alternans was derived from 32 consecutive beats using a spectral method as previously described [20, 21], where the k score was >3 and the alternans magnitude (Valt) was >0.02 mV. The rate threshold for alternans was taken as the longest paced cycle length to show alternans. A dynamic restitution curve was created by plotting paced cycle length against APD90 and fitting a single exponential curve using Equation 1, where a and b are real number variables. The gradient of the slope at each paced cycle length was calculated using Equation 2 and the rate at which the gradient became >1 was noted (the point at which action potential alternans is significantly more likely to occur [22]).

Equation 1:

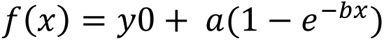

Equation 2:

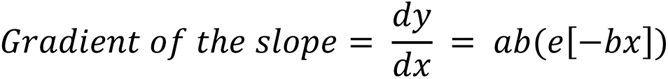

Atrial conduction velocity was recorded using a 2/8/2 mm spaced decapolar catheter (Biosense Webster, Irvine, CA) placed parallel to the lateral wall of the right atrium. Pacing was performed across the distal bipoles (bipoles 1-2) via a Medtronic Carelink 2090 programmer over a cycle lengths range of 500 to 270 ms. Signals were digitised at 1kHz using a PowerLab amplifier and analysed in LabChart, where conduction velocity was calculated the distance (based on the fixed interelectrode spacing) and difference in bipolar electrogram timing on bipoles 3-4 versus bipoles 7-8.

Vulnerability to induced AF was assessed by delivering 50 Hz bursts at 5 V for 5 s via the ICD to the right atrium. Arrhythmia burden was measured as the duration of AF upon cessation of high frequency stimulation until restoration of sinus rhythm averaged over ten inductions. A cut-off of 15 minutes duration was pre-specified, after which AF was terminated either by overdrive pacing or cardioversion through the ICD. The dominant frequency of induced AF was calculated when a minimum threshold of 5 s of fibrillation was present [23]. After bandpass filtering between 3-15 Hz, a Hamming data window was applied and a power spectrum created in LabChart with a frequency resolution of 0.25 Hz.

Atrial wavelength and fibrillation number were calculated as previously described [24, 25], where wavelength is the product of conduction velocity and AERP and fibrillation number is the left atrial diameter (on echo) divided by the wavelength.

### Administration of sildenafil

Upon completion of the baseline assessment and electrophysiology study, 10 mg of sildenafil (Revatio; Pzifer, USA) was administered as an intravenous bolus. After a five-minute stabilization period, all electrophysiological experiments were repeated. Upon study completion, sheep were euthanized with 200 mg.kg^-1^ intravenous pentobarbitone.

### Data presentation and statistics

Data are reported as mean ± standard error of the mean (SEM). Non-normalised datasets, as determined by Shapiro-wilk tests, were log-transformed to achieve normality where appropriate. If any values equalled zero, then non-parametric tests were used instead of a transform. A two-tailed student’s T-test and Fisher’s exact test were used to compare continuous and categorical data respectively in GraphPad Prism v10 (GraphPad Software, San Diego, CA). Differences between non-linear curves was assessed using the sum-of-squares F test, where separation at the 95% confidence intervals was taken as significant. Statistical significance was taken as a p value <0.05. Exact P values are reported.

## Results

### Sildenafil reduced the vulnerability to induced AF

All twelve sheep completed electrophysiology studies. (**Figure 1A)**. Sildenafil decreased the mean duration of AF over the ten inductions per sheep (112.2 ± 73.5 s vs. 3.3 ± 1.4 s; *p*=0.007; **Figure 1B**). The decrease in AF with sildenafil was due to a decrease in the number of inductions that resulted in AF (90 % vs 70 % in controls vs. sildenafil respectively; *p*=0.001; **Figure 1C**) and a tendency to shorter episodes, when AF occurred, although this did not reach statistical significance (117.0 ± 73.4 s vs. 8.3 ± 3.3 s; *p*=0.09; **Figure 1D**). The dominant frequency of induced AF was decreased after sildenafil administration (5.5 ± 0.2 Hz vs 4.8 ± 0.3 Hz; *p*=0.012; **Figure 1E**).

**Figure 1.**
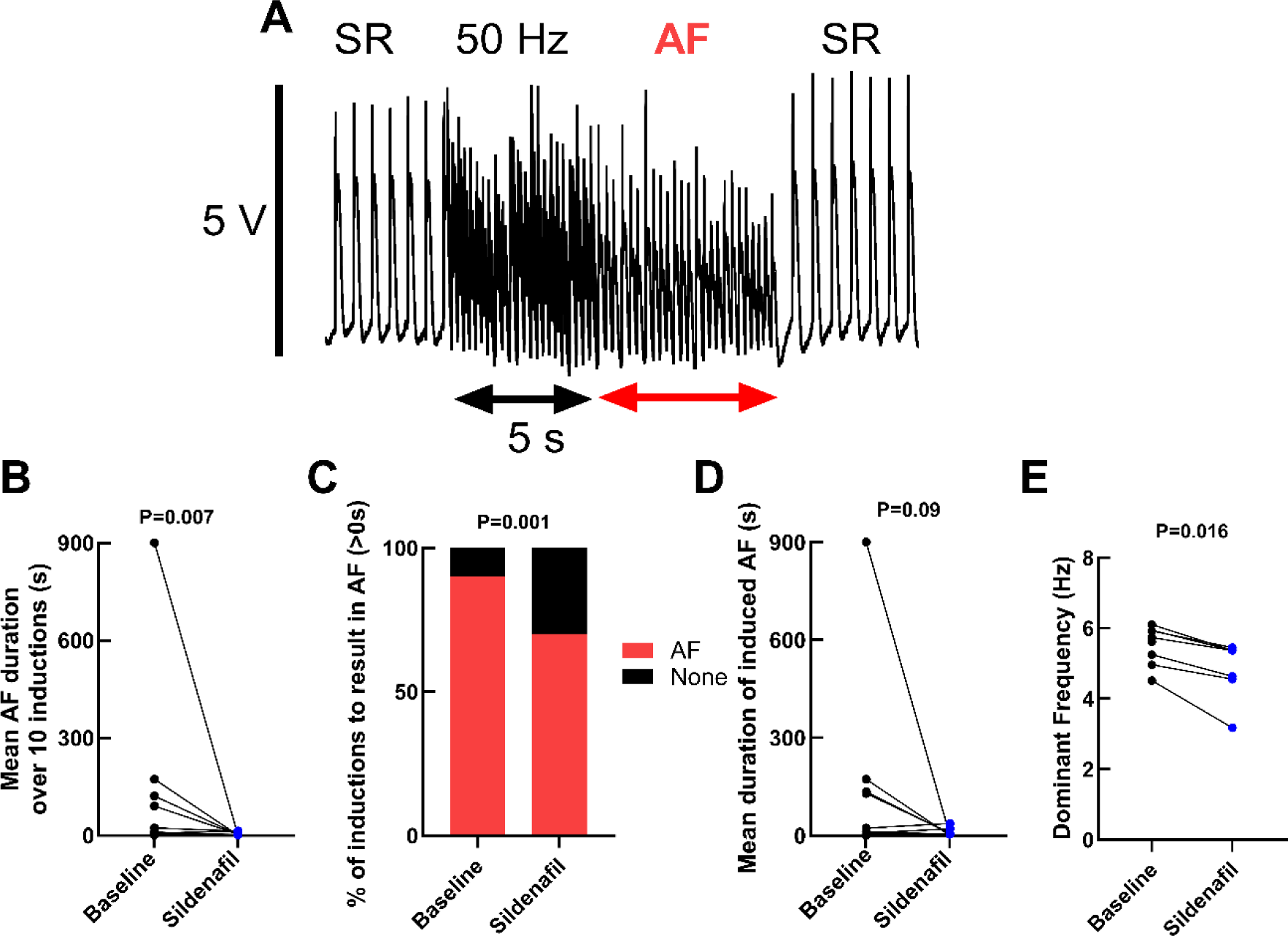
Sildenafil reduces the burden of induced AF. (A) Example trace showing AF duration was recorded after each five second, 50 Hz burst from cessation of the burst until spontaneous reversion to sinus rhythm. (B) sildenafil reduced the mean duration of AF over ten inductions. (C) sildenafil resulted in fewer inductions leading to AF. (D) sildenafil did not reduce the duration of AF when induced. (E) sildenafil reduced dominant frequency of induced AF. N=12.

### Sildenafil had no effect on baseline electrocardiographical parameters

Sildenafil did not affect ECG parameters including; PR interval (111 ± 3 ms vs. 115 ± 4 ms, p=0.38) and PWD_max_ (53 ± 1 ms vs. 50 ± 1 ms, p=0.08), or QRS duration (50 ± 2 ms vs. 52 ± 1 ms, p=0.17) and the uncorrected QT interval (336 ± 12 ms vs. 337 ± 10 ms, p=0.88).

Sildenafil reduced LVSP (94 ± 7 mmHg vs. 75 ± 4 mmHg, p=0.01) however had no effect on LVEDP (14 ± 3 mmHg vs. 10 ± 1 mmHg, p=0.2). There was an increase in heart rate associated with sildenafil (cycle length: 854 ± 46 ms vs. 639 ± 17 ms, p=0.01; **Table 1**) likely as a reflex response to the fall in LVSP.

**Table 1.** Sildenafil does not alter ECG parameters. A 5-lead surface ECG recording of changes pre- and post-sildenafil was performed in ten sheep and invasive blood pressure measurements were recorded from the left ventricle in six sheep. PWD_Max_ is maximal P wave duration, LVSP is left ventricular systolic pressure and LVEDP is left ventricular end diastolic pressure.

|  | **Baseline** | **Sildenafil** | **P value** |
| --- | --- | --- | --- |
| **PR interval (ms)** | 111 ± 3 | 115 ± 4 | 0.38 |
| **PWD<sub>Max</sub> (ms)** | 53 ± 1 | 50 ± 1 | 0.08 |
| **Cycle length (ms)** | 854 ± 46 | 639 ± 17 | 0.0002 |
| **QRS duration**<br>**(ms)** | 50 ± 2 | 52 ± 1 | 0.17 |
| **QT (ms)** | 336 ± 12 | 337 ± 10 | 0.88 |
| **LVSP (mmHg)** | 94 ± 7 | 75 ± 4 | 0.01 |
| **LVEDP (mmHg)** | 14 ± 3 | 10 ± 1 | 0.2 |

### Sildenafil prolonged the atrial effective refractory period

To determine sildenafil’s anti-arrhythmic mechanism, we measured the electrophysiological properties of the right atrium. Sildenafil prolonged the AERP by approximately 20 ms (146.9 ± 7.2 ms vs 166.2 ± 32.5 ms; p=0.006; **Figures 2A-B**), however it had no effect on atrial conduction velocity over the range of paced cycle lengths, with an average velocity of 0.88 ± 0.003 m.s^-1^ in both conditions (p=0.99, **Figures 2C-D**). As atrial wavelength is the product of AERP and conduction velocity, the prolongation of the AERP increased the mean wavelength in sildenafil (12.9 ± 0.07 cm vs 15.0 ± 0.07 cm; P<0.0001; **Figure 2E**). The wavelength was divided by atrial diameter for each animal (2.5 ± 0.08 cm; **Figure 2F-G**) to derive the fibrillation number as an index of AF vulnerability. Sildenafil decreased fibrillation number by approximately 10% due to the increased atrial wavelength (0.20 ± 0.001 vs 0.18 ± 0.001; P<0.0001; **Figure 2H**).

**Figure 2.**
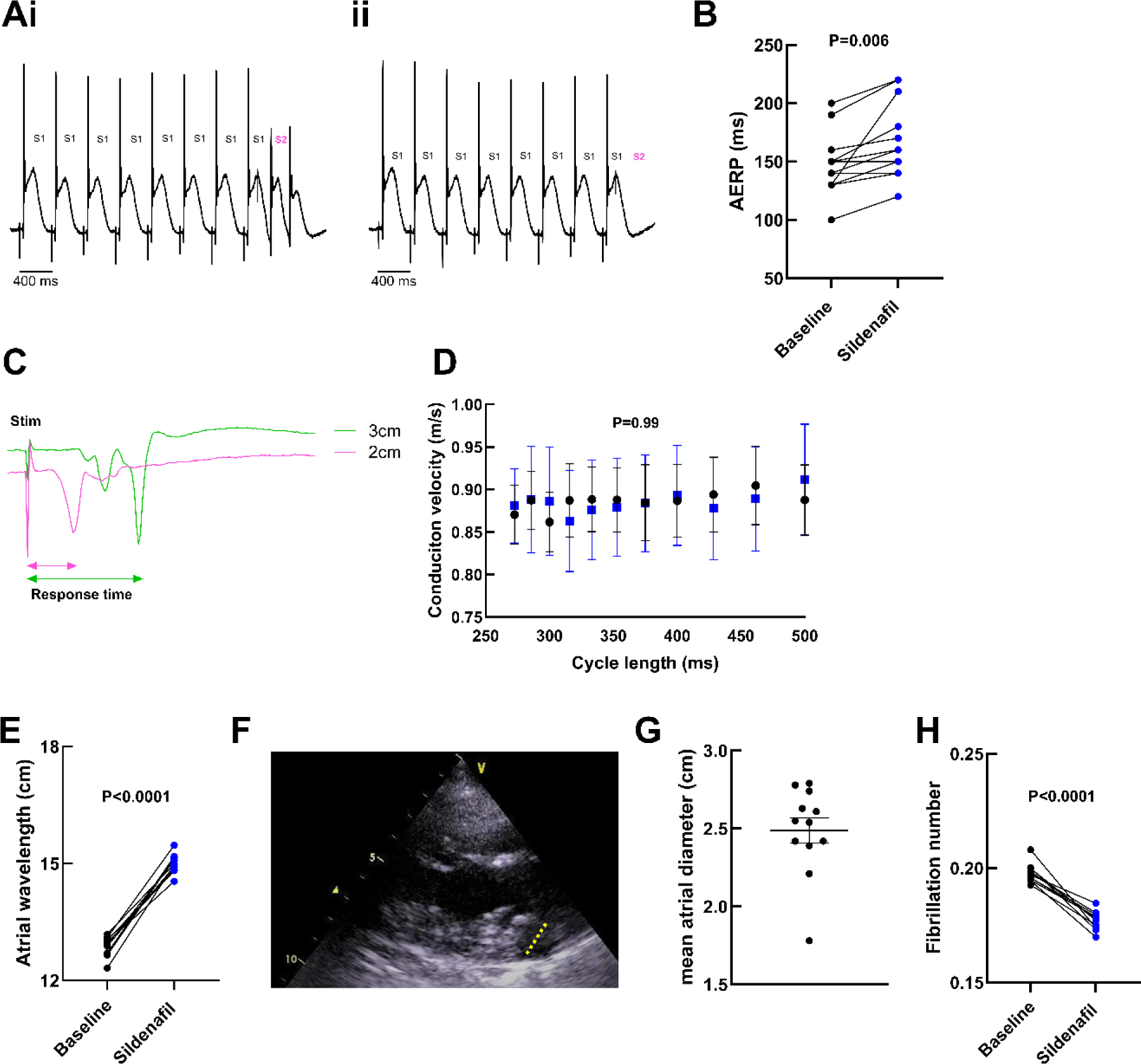
Atrial electrophysiological changes in sildenafil are predicted to be anti-arrhythmic. (A) Example traces showing a programmed electrical stimulation study of eight S1 stimulations with a 400 ms drive cycle length followed by an S2 (pink) decremented in 10 ms decrements each cycle that elicited (i) and failed to elicit (ii) a MAP. (B) AERP, calculated as the longest cycle length at of which the S2 stimulation failed to elicit a MAP. (C) Example traces showing calculation of conduction velocity from the mean time lag between stimulation and response at biploar electrodes spaced 2cm (pink) and 3cm (green) from the stimulating catheter. (D) conduction velocity calculated from 500 ms to 250 ms cycle lengths. (E) Atrial wavelength calculated as the product of the AERP and the average conduction velocity. (F) Left atrial diameter measured over three cycles in the anteroposterior axis (yellow dotted line) whilst the sheep was conscious. (G) Atrial diameter. (H) Fibrillation number calculated from wavelength divided by atrial diameter pre and post sildenafil as an index of AF vulnerability. N=11.

### AERP prolongation is not due to a change in action potential duration

The main electrophysiological mechanism by which Sildenafil reduced the vulnerability to induced AF appeared to be through prolongation of the AERP rather than a change in atrial conduction velocity. We next investigated whether this was due to a prolongation of the MAP. The steady state APD90 for MAPs was calculated over a range of paced cycle lengths as shown in **Figure 3A**. Sildenafil particularly prolonged the APD90 at lower cycle lengths (Cycle length of 500ms: 181.2 ± 8.7 vs 215.2 ± 17.0, p=0.02 (paired t-test), overall p=0.06; **Figure 3B**).

**Figure 3.**
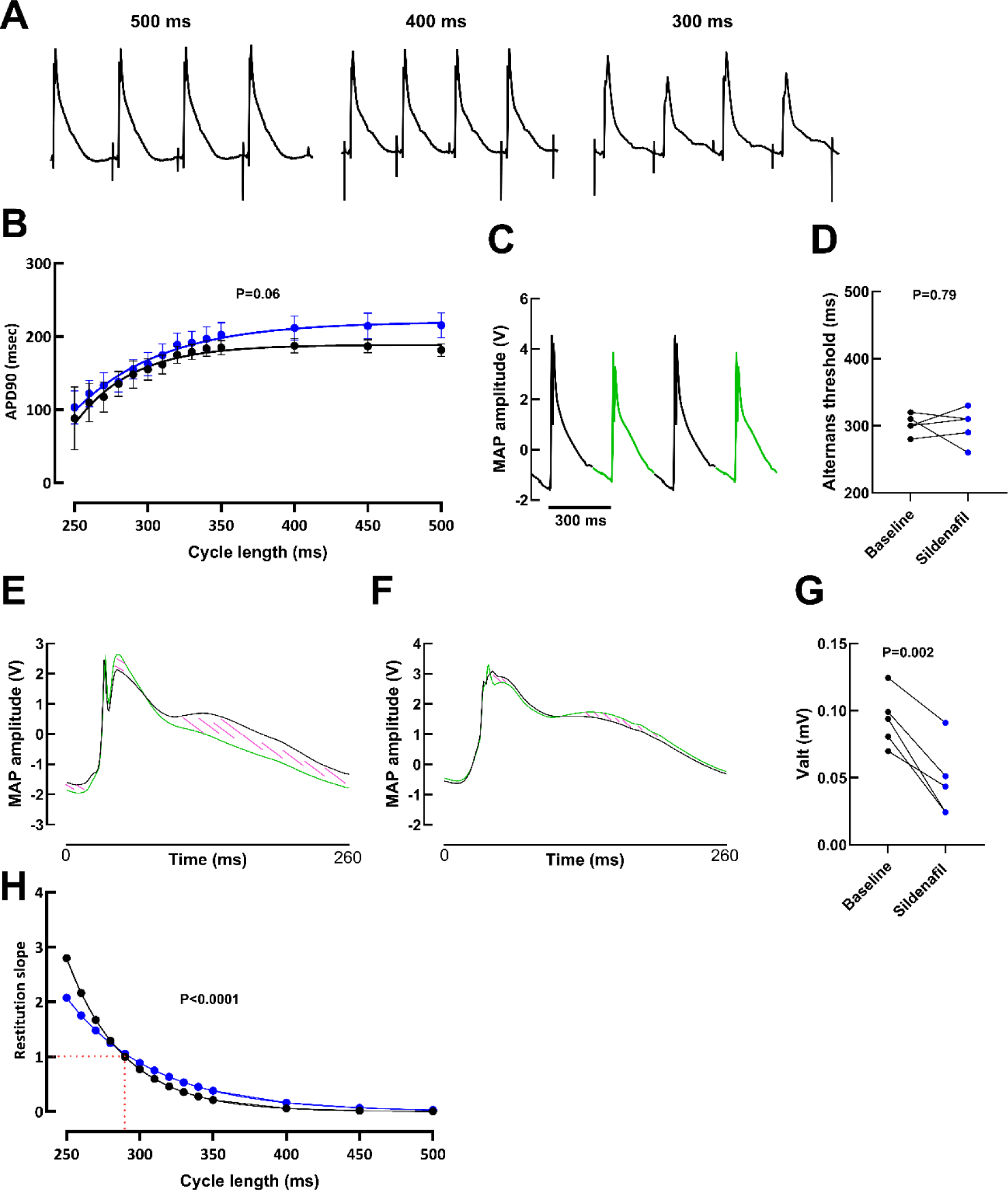
Sildenafil shallows MAP restitution and alternans magnitude. (A) Example MAPs at 500 ms, 400 ms and 300 ms. (B) APD90. calculated from 32 consecutive MAPs at each cycle length, plotted against cycle length and non-linear first order exponential decay curves fitted. (C) Example trace showing alternating odd (black) and even (green) MAPs. (D) Threshold for alternans calculated as the slowest rate at which alternans could be detected using a discrete Fourier transform and established parameters of K score > 3 and Valt > 0.02 mV. (E) Alternating MAPs overlaid and the integrated area difference between MAPs (shaded pink) calculated as the magnitude of alternans (Valt) pre-sildenafil and (F) post-sildenafil at 260 ms cycle length stimulation in the same animal. (G) Magnitude of alternans (Valt) at threshold. (H) Mean gradient of the rate of APD90 change between cycle lengths calculated from differentiation of the curves which is indicative of a role for alternans. N=11.

### Sildenafil had no effect on the threshold for atrial alternans

Atrial alternans has been linked with the initiation of AF in patients and animal models and therefore we next investigated if Sildenafil altered the propensity for alternans. The rate threshold for atrial MAP alternans was calculated before and after Sildenafil administration in 6 sheep in which alternans threshold could be determined, the other 6 did not exhibit alternans by a cycle length of 250 ms. Sildenafil did not affect the rate threshold for these sheep (301.7 ± 5.4 ms vs 305.0 ± 10.9 ms; p=0.79; **Figure 3C and D**), however the alternans amplitude, Valt was decreased by approximately 50 % in the presence of Sildenafil (0.09 ± 0.001 mV vs. 0.05 ± 0.10 mV; p=0.002; **Figure 3E, F and G**). Given we could only calculate the atrial MAP alternans threshold in six of the twelve sheep, we examined the gradient of the restitution curve as a second marker of arrhythmogenicity. Sildenafil reduced the gradient of the curve (K=0.03 vs 0.02; p<0.0001; **Figure 3H**) suggesting it may have an enhanced antiarrhythmic effect at faster atrial rates shorter than 280 ms where the gradient of the curve is >1, indicated by the red dotted line.

## Discussion

The key findings of this study were, i) sildenafil decreased AF burden by decreasing the AF complexity, the number of AF episodes and tending to decrease AF duration, ii) sildenafil increased the AERP which increased the excitation wavelength and likely contributes to the tendency for sildenafil to decrease AF duration, iii) sildenafil decreased the slope of the AP restitution curve which was associated with a decrease in the amplitude of alternans and could reduce the propensity for AF genesis.

### Acute sildenafil treatment reduces vulnerability to AF

Our data suggests that acute sildenafil has anti-arrhythmic properties which reduce the vulnerability to induced AF. No previous studies have directly examined the effects of PDE5 inhibition on atrial arrhythmias however intermittent sildenafil has previously been associated with a reduced incidence of AF in a retrospective observational study in type II diabetic males [1]. Treating diabetes would be predicted to reduce AF incidence but our data suggests this effect was due to the direct effect of sildenafil. This study did not specify the amount of sildenafil taken, nor the degree of arrhythmia monitoring, therefore a cause-and-effect relationship between sildenafil and AF could not be established. Interestingly, a number of case reports have been published which associate sildenafil use with the acute onset of AF [26–28]. These reports are characterized initially by a state of profound hypotension followed by a reflex tachycardia which suggests AF was induced in a state of catecholamine excess. Sympathetic stimulation is recognized as a cause of AF particularly in those with pre-existing structural heart disease [29, 30]. A mean fall in systolic blood pressure and a rise in heart rate were observed in this study which may have resulted from a change in autonomic tone or systemic vasodilation associated with PDE5 inhibition within vascular smooth muscle. Sympathetic stimulation with isoprenaline has previously been reported to show pro-arrhythmic effects such as shortening of the AERP and increasing the inducibility of AF [31, 32]. Our findings suggest the mechanism of action of sildenafil is not solely dependent on inducing changes in autonomic activity.

Acute sildenafil has been studied for its anti-arrhythmic properties in the ventricles. We have previously demonstrated the anti-arrhythmic properties of acute sildenafil in a sheep model of drug induced long QT which was found to be resultant of a reduced number of premature ventricular complexes and afterdepolarizations, and fewer R on T events due to a decreased SR calcium content [5]. Similarly, canine study found that acute sildenafil reduced the vulnerability to ventricular arrhythmias in the setting of acute myocardial ischemia [4]. Sildenafil reduced ventricular ectopy, tachycardia and fibrillation within the first 24 hours however no electrophysiology study was performed, and the underlying electrophysiological mechanism is therefore unclear.

### Acute sildenafil treatment reduces alternans magnitude

Alternans is thought to be both an important substrate and trigger of AF where alternans has been seen prior to AF and correlates with AF inducibility [21, 33, 34], so we sought to determine whether alternans was affected by acute sildenafil. There was no difference in the rate threshold for atrial alternans after sildenafil. This was despite the dynamic restitution curve showing sildenafil reduced the maximum gradient, which would ordinarily be expected to decrease the probability of developing alternans. It is recognized however that anti-arrhythmic drugs which alter the slope of the restitution curve alone are unlikely to be successful unless they also have effects on regional heterogeneity on electrophysiological parameters (APD90 and AERP for example) and conduction velocity restitution [35]. This may potentially explain why changing the maximum gradient on curve did not impact on the alternans threshold. A steeper restitution gradient is indicative of a greater magnitude of alternans [36]. Sildenafil did halve the magnitude of alternans at threshold. The magnitude of alternans has been shown to be important in AF genesis, where the magnitude of alternans has been seen to mirror the progression of AF and is seen to be greatest just before transitioning into AF [37, 38]. Our data therefore suggests that by decreasing the amplitude of MAP alternans, sildenafil decreases the likelihood alternans will trigger AF physiologically.

### The main anti-arrhythmic effect of sildenafil is on the AERP

Sildenafil prolonged the AERP which also resulted in an increase in the wavelength. The multiple wavelet hypothesis of AF states that multiple re-entrant waves of excitation are being generated throughout the atria and are then propagated to adjacent excitable tissue [39]. AF is sustained as although these wavelets are extinguished when they either collide with each other or unexcitable tissue, enough new wavelets are being created to maintain the arrhythmia. This hypothesis also predicts AF will terminate when all wavelets are extinguished. The anatomy of a single wavelet has been refined over the years from the leading circle model to the spiral wave [40] however the wavelength (the product of ERP and conduction velocity) remains a core feature.

An increase in the AERP by sildenafil without a change in conduction velocity drove an increase in the wavelength. Conceptually, this means sildenafil changes the electrophysiological properties of the atria such that fewer re-entrant wavelets can be accommodated in a fixed volume of atrial tissue therefore there is a greater probability they will all extinguish at a single time-point and terminate AF. Sildenafil also reduced the dominant frequency of induced AF, which represents the activity of a “mother rotor” (a single dominant wavelet which maintains all other re-entrant wavelets in the atria). A lower dominant frequency corresponds to a rotor with a slower rotation time, a larger core size and decreased curvature, all of which are considered anti-arrhythmic as fewer larger spiral waves may be accommodated in a fixed volume of tissue. The anti-arrhythmic actions of sildenafil on AERP, wavelength and dominant frequency suggests these are important mechanisms reducing the vulnerability to AF. The ionic mechanism behind the prolongation of AERP was not investigated in this study but it is known that the AERP is determined by the APD90, which was prolonged by sildenafil, predominantly at slower pacing rates.

Assumptions have also been made that the anti-arrhythmic effects are solely due to PDE5 inhibition effecting the cGMP-PKG axis. There is the possibility of crosstalk where higher levels of cGMP lead to competition for PDE3, enhancing the cAMP-PKA axis. The evidence base in the literature would not support this mechanism as increasing PKA activity by crosstalk has been reported to increase ventricular systolic pressure [42] whereas the opposite occurred in this study. Crosstalk with the PDE2 compartment is also possible however to our knowledge, there are no studies looking at specific PDE2 inhibition and the vulnerability to AF.

### Limitations

The electrophysiology study was performed at a single site in the right atrium only. Regional differences in APD and AERP are known to occur due to changes in ion channel expression [43–45]. It was not possible to measure in the LA using an endocardial approach due to difficulties in access. Despite this, there is no reason to believe the anti-arrhythmic effects seen would have been specific to the right atrium only. Repeated burst pacing is often used as a mechanism by which to induce permanent AF [24, 46], and repeating alternans inductions is known to increase propensity to alternans [47]. As we have performed a baseline study and then repeated this in the presence of sildenafil, rather than comparing to time controls, the true reduction of AF burden in sildenafil and alternans magnitude and possibly alternans threshold may be obfuscated by our approach and actually greater than observed.

Finally, the study was performed acutely in healthy sheep in the absence of any pre-existing atrial remodeling. Although anti-arrhythmic effects of PDE5 inhibition have been observed, it is unclear whether these effects will also be seen in chronic models of AF or in humans with AF which warrants investigation in the future.

## Conclusion

Sildenafil administration drastically reduces AF vulnerability in healthy sheep atria through a prolonged AERP and the wavelength, reduction of the dominant frequency of the AF rotors, a shallowing of the restitution curve and a smaller magnitude of alternans. Whether sildenafil could be used to treat or prevent AF in a clinical setting is yet to be determined but is an important avenue to explore.

## Author contributions

ND, GM, DC, CS, AW, MO, CP animal model and surgical EP studies. ND, GM analysis and interpretation, AW, CP, KD conception, funding, supervision. All authors involved in drafting and are in agreement with final manuscript.

## References

1. Anderson, S.G., et al., Phosphodiesterase type-5 inhibitor use in type 2 diabetes is associated with a reduction in all-cause mortality. Heart, 2016. 102(21): p. 1750–1756.

2. Lin, Y., et al., Various subtypes of phosphodiesterase inhibitors differentially regulate pulmonary vein and sinoatrial node electrical activities. Experimental and Therapeutic Medicine, 2020. 19(4): p. 2773–2782.

3. Lee, T., et al., Effect of sildenafil on ventricular arrhythmias in post-infarcted rat hearts. European Journal of Pharmacology, 2012. 690(1-3): p. 124–132.

4. Nagy, O., et al., Sildenafil (Viagra) reduces arrhythmia severity during ischaemia 24 h after oral administration in dogs. Br J Pharmacol, 2004. 141(4): p. 549–51.

5. Hutchings, D., et al., PDE5 Inhibition Suppresses Ventricular Arrhythmias by Reducing SR Ca2+ Content. Circulation Research, 2021. 129(6): p. 650–665.

6. Haddad, G.E., N. Sperelakis, and G. Bkaily, Regulation of the calcium slow channel by cyclic GMP dependent protein kinase in chick heart cells. Mol Cell Biochem, 1995. 148(1): p. 89–94.

7. Abi-Gerges, N., R. Fischmeister, and P.F. Mery, G protein-mediated inhibitory effect of a nitric oxide donor on the L-type Ca2+ current in rat ventricular myocytes. J Physiol, 2001. 531(Pt 1): p. 117–30.

8. Simmons, M.A. and H.C. Hartzell, Role of phosphodiesterase in regulation of calcium current in isolated cardiac myocytes. Mol Pharmacol, 1988. 33(6): p. 664–71.

9. Campbell, D.L., J.S. Stamler, and H.C. Strauss, Redox modulation of L-type calcium channels in ferret ventricular myocytes. Dual mechanism regulation by nitric oxide and S-nitrosothiols. J Gen Physiol, 1996. 108(4): p. 277–93.

10. Wahler, G.M. and N. Sperelakis, Intracellular injection of cyclic GMP depresses cardiac slow action potentials. J Cyclic Nucleotide Protein Phosphor Res, 1985. 10(1): p. 83–95.

11. Mery, P.F., et al., Ca2+ current is regulated by cyclic GMP-dependent protein kinase in mammalian cardiac myocytes. Proc Natl Acad Sci U S A, 1991. 88(4): p. 1197–201.

12. Levi, R.C., G. Alloatti, and R. Fischmeister, Cyclic GMP regulates the Ca-channel current in guinea pig ventricular myocytes. Pflugers Arch, 1989. 413(6): p. 685–7.

13. Fischmeister, R. and H.C. Hartzell, Cyclic guanosine 3’,5’-monophosphate regulates the calcium current in single cells from frog ventricle. J Physiol, 1987. 387: p. 453–72.

14. Hartzell, H.C. and R. Fischmeister, Opposite effects of cyclic GMP and cyclic AMP on Ca2+ current in single heart cells. Nature, 1986. 323(6085): p. 273–5.

15. Wang, H., et al., Phosphodiesterase 5 restricts NOS3/Soluble guanylate cyclase signaling to L-type Ca2+ current in cardiac myocytes. J Mol Cell Cardiol, 2009. 47(2): p. 304–14.

16. Vandecasteele, G., et al., Cyclic GMP regulation of the L-type Ca(2+) channel current in human atrial myocytes. J Physiol, 2001. 533(Pt 2): p. 329–40.

17. Nagayama, T., et al., Sildenafil stops progressive chamber, cellular, and molecular remodeling and improves calcium handling and function in hearts with pre-existing advanced hypertrophy caused by pressure overload. J Am Coll Cardiol, 2009. 53(2): p. 207–15.

18. Gong, W., et al., Chronic inhibition of cGMP-specific phosphodiesterase 5 suppresses endoplasmic reticulum stress in heart failure. Br J Pharmacol, 2013. 170(7): p. 1396–409.

19. Chesnais, J.M., R. Fischmeister, and P.F. Mery, Positive and negative inotropic effects of NO donors in atrial and ventricular fibres of the frog heart. J Physiol, 1999. 518 **( Pt** **2**): p. 449–61.

20. Pearman, C.M., An Excel-based implementation of the spectral method of action potential alternans analysis. Physiol Rep, 2014. 2(12).

21. Pearman, C., et al., Increased Vulnerability to Atrial Fibrillation Is Associated With Increased Susceptibility to Alternans in Old Sheep. Journal of the American Heart Association, 2018. 7(23).

22. Pruvot, E.J., et al., Role of calcium cycling versus restitution in the mechanism of repolarization alternans. Circ Res, 2004. 94(8): p. 1083–90.

23. Stiles, M.K., et al., The effect of electrogram duration on quantification of complex fractionated atrial electrograms and dominant frequency. J Cardiovasc Electrophysiol, 2008. 19(3): p. 252–8.

24. Denham, N., et al., Optimising Large Animal Models of Sustained Atrial Fibrillation: Relevance of the Critical Mass Hypothesis. Frontiers in Physiology, 2021. 12.

25. Hwang, M., et al., Fibrillation number based on wavelength and critical mass in patients who underwent radiofrequency catheter ablation for atrial fibrillation. IEEE Trans Biomed Eng, 2015. 62(2): p. 673–9.

26. Hahn, I.H. and R.S. Hoffman, Aroused to atrial fibrillation? Am J Emerg Med, 2000. 18(5): p. 642.

27. Awan, G.M., et al., Acute, symptomatic atrial fibrillation after sildenafil citrate therapy in a patient with hypertrophic obstructive cardiomyopathy. Am J Med Sci, 2000. 320(1): p. 69–71.

28. Hayashi, K., et al., Atrial fibrillation and continuous hypotension induced by sildenafil in an intermittent WPW syndrome patient. Jpn Heart J, 1999. 40(6): p. 827–30.

29. Huang, J.L., et al., Changes of autonomic tone before the onset of paroxysmal atrial fibrillation. Int J Cardiol, 1998. 66(3): p. 275–83.

30. Dimmer, C., et al., Variations of autonomic tone preceding onset of atrial fibrillation after coronary artery bypass grafting. Am J Cardiol, 1998. 82(1): p. 22–5.

31. Krummen, D.E., et al., Mechanisms of human atrial fibrillation initiation: clinical and computational studies of repolarization restitution and activation latency. Circ Arrhythm Electrophysiol, 2012. 5(6): p. 1149–59.

32. Farges, J.P., M. Ollagnier, and G. Faucon, Influence of acetylcholine, isoproterenol, quinidine and ouabain on effective refractory periods of atrial and ventricular myocardium in the dog. Arch Int Pharmacodyn Ther, 1977. 227(2): p. 206–19.

33. Narayan, S.M., et al., Repolarization alternans reveals vulnerability to human atrial fibrillation. Circulation, 2011. 123(25): p. 2922–30.

34. Narayan, S.M., et al., Alternans of atrial action potentials during atrial flutter as a precursor to atrial fibrillation. Circulation, 2002. 106(15): p. 1968–73.

35. Banville, I. and R.A. Gray, Effect of action potential duration and conduction velocity restitution and their spatial dispersion on alternans and the stability of arrhythmias. J Cardiovasc Electrophysiol, 2002. 13(11): p. 1141–9.

36. Koller, M., et al., Altered dynamics of action potential restitution and alternans in humans with structural heart disease. Circulation, 2005. 112(11): p. 1542–1548.

37. Lalani, G., et al., Frequency Analysis of Atrial Action Potential Alternans A Sensitive Clinical Index of Individual Propensity to Atrial Fibrillation. Circulation-Arrhythmia and Electrophysiology, 2013. 6(5): p. 859–867.

38. Monigatti-Tenkorang, J., et al., Intermittent atrial tachycardia promotes repolarization alternans and conduction slowing during rapid rates, and increases susceptibility to atrial fibrillation in a free-behaving sheep model. J Cardiovasc Electrophysiol, 2014. 25(4): p. 418–27.

39. Moe, G.K., W.C. Rheinboldt, and J.A. Abildskov, A COMPUTER MODEL OF ATRIAL FIBRILLATION. Am Heart J, 1964. 67: p. 200–20.

40. Comtois, P., J. Kneller, and S. Nattel, Of circles and spirals: bridging the gap between the leading circle and spiral wave concepts of cardiac reentry. Europace, 2005. 7 **Suppl 2**: p. 10–20.

41. Franz, M.R., et al., Drug-induced post-repolarization refractoriness as an antiarrhythmic principle and its underlying mechanism. Europace, 2014. 16 **Suppl 4**: p. iv39–iv45.

42. Nagendran, J., et al., Phosphodiesterase type 5 is highly expressed in the hypertrophied human right ventricle, and acute inhibition of phosphodiesterase type 5 improves contractility. Circulation, 2007. 116(3): p. 238–48.

43. Li, D., et al., Potential ionic mechanism for repolarization differences between canine right and left atrium. Circ Res, 2001. 88(11): p. 1168–75.

44. Al Abed, A., N.H. Lovell, and S. Dokos, Local Heterogeneous Electrical Restitution Properties of Rabbit Atria. J Cardiovasc Electrophysiol, 2016. 27(6): p. 743–53.

45. Feng, J., et al., Ionic mechanisms of regional action potential heterogeneity in the canine right atrium. Circ Res, 1998. 83(5): p. 541–51.

46. Enriquez-Vazquez, D., et al., Non-invasive electromechanical assessment during atrial fibrillation identifies underlying atrial myopathy alterations with early prognostic value. Nature Communications, 2023. 14(1).

47. Walker, M., et al., Hysteresis effect implicates calcium cycling as a mechanism of repolarization alternans. Circulation, 2003. 108(21): p. 2704–2709.

